# Homogeneous antibody-DNA-conjugates using unmodified oligonucleotides and photo-crosslinkable protein G-HUH endonuclease fusion proteins

**DOI:** 10.1101/2025.11.14.688079

**Authors:** Anna Swietlikowska, Femi Hesen, Alexander Gräwe, Maarten Merkx

## Abstract

Antibody-DNA conjugates are increasingly used in analytical biochemistry and nanotechnology. However, current methods for their preparation often lack specificity, resulting in heterogeneous products. Here, we present a novel strategy to covalently label the Fc domain of antibodies with single-strand DNA in a site-specific and stoichiometrically controlled manner. This method employs unmodified oligonucleotides and a fusion protein consisting of a protein G dimer linked to an HUH endonuclease. We first evaluated the sequence specificity and optimal conditions for DNA attachment to three HUH endonuclease variants, DCV, PCV2, and WDV. Our results show increased sequence specificity in the presence of Mg^2+^ compared to the more commonly used Mn^2+^ cofactor ion. Although formation of the phospho-tyrosine bond is found to be reversible, no significant hydrolysis of the protein-DNA conjugates is observed for up to 8 days at room temperature. The DCV domain allowed essentially complete formation of DNA-fusion protein conjugates at a 1:1 protein-DNA ratio, eliminating the need for removal of excess oligos. Subsequent photo-crosslinking yielded antibody-DNA conjugates that were successfully used in a proximity extension assay (PEA) to detect TNFα and IL-6. Our method provides an efficient, low cost and generally applicable method to prepare homogeneous antibody–DNA conjugates with 1:1 stoichiometry.

## Introduction

Antibody-DNA conjugates are versatile macromolecules used in applications such as immuno-PCR for protein detection,^1–4^ DNA-PAINT for cell immunophenotyping^5,6^ and the functionalisation of DNA Origami nanostructures.^7,8^ Typically, DNA is covalently linked to antibodies through reactive lysine or cysteine residues using oligonucleotides modified with chemical groups such as NHS-esters or maleimides.^9^ However, these approaches offer limited control over both the conjugation site and the resulting stoichiometry. Covalent modifications also require additional purification steps to remove excess reactants, decreasing the yield of antibody conjugates. The efficiency of the maleimide-cysteine reaction can also decrease with increasing DNA length, limiting its compatibility with longer sequences such as DNA aptamers.^10^ While targeting glycan groups on the Fc domain can provide site specificity, controlling the stoichiometry of this reaction remains challenging.^11,12^ Moreover, these strategies rely on expensive modified oligonucleotides, multi-step protocols and specialized expertise.

To address some of these limitations, we and others previously reported the use of photo-crosslinkable protein G-DNA conjugates to enable site-specific attachment of oligonucleotides. ^13–15^ This approach is based on the LASIC method^16^ in which protein G is engineered to incorporate the unnatural amino acid pBPA (p-benzoylphenylalanine). Upon UV irradiation, pBPA forms a covalent bond with a methionine residue in the Fc domain of mammalian IgG antibodies, enabling stable conjugation. However, this method still requires a 2-step reaction in which single-strand DNA (ssDNA) with a 5’-amine group is first reacted with the bifunctional cross-linker sulfo-SMCC, followed by reaction of the maleimide-functionalized DNA with a cysteine introduced in protein G.

Endonucleases from the HUH family have recently been introduced as efficient protein tags that enable sequence-specific, covalent attachment of unmodified ssDNA to proteins.^17–22^ Their use alleviates the need for modified DNA and substantially accelerates DNA conjugation. The conserved HUH motif, which consists of two histidines separated by a hydrophobic residue, coordinates a divalent ion cofactor (magnesium or manganese) which coordinates the phosphate ester backbone and activates it for attack by a catalytic tyrosine, forming a 5’-phosphotyrosine covalent bond.^23^ Within the HUH superfamily, proteins of the Rep (replication initiator protein) subgroup are particularly attractive for protein-DNA conjugation due to their small size (∼11-16 kDa) and short recognition sequence (∼9-nt), which has enabled applications in genome editing,^18^ cell targeting,^20^ and biosensing.^19^ The molecular basis of DNA recognition by the Rep domains has been elucidated by the Gordon group through a comprehensive exploration of their sequence specificity and by solving the crystal structure of several Reps, including DCV, PCV2 and WDV. ^22,24–27^

In this study, we combine photo-crosslinkable protein G with HUH tag-mediated DNA labelling to develop a site-specific, stoichiometrically-controlled method for antibody-DNA conjugation using unmodified DNA oligonucleotides. We designed and characterized 3 fusion proteins, each comprising of a photo-crosslinkable protein G dimer (pG) fused to different endonuclease domains: DCV, PCV2 and WDV. Building on previous work by the Gordon group, we examined the influence of manganese and magnesium ions on sequence specificity, and further assessed the pH dependence, stability, and reversibility of the conjugation reaction. To demonstrate functionality, antibody-DNA conjugates were generated through photo-crosslinking of DNA-functionalized pG-DCV fusion proteins (Figure 1), followed by their application in proximity extension immunoassays (PEA).^2,3^

**Figure 1.**
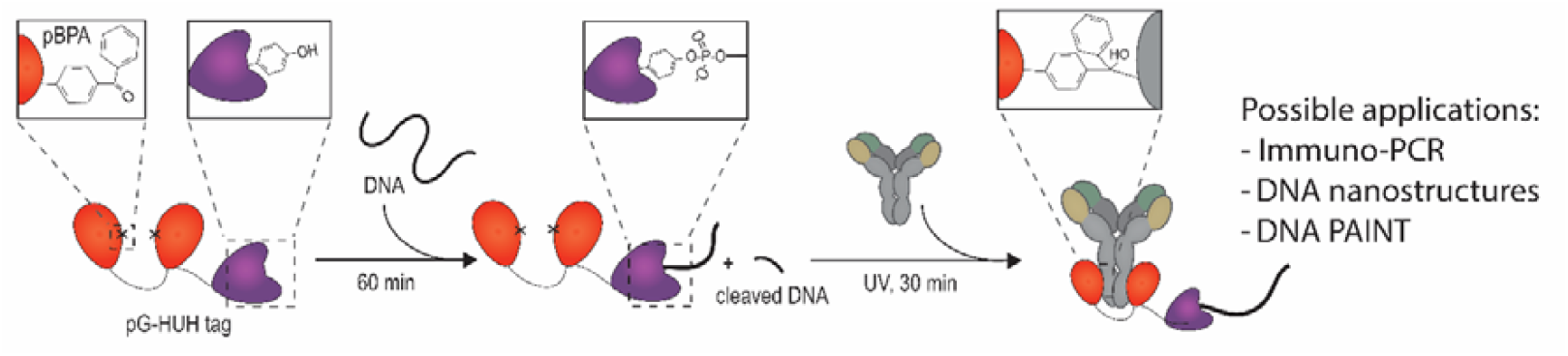
Efficient antibody labelling with unmodified DNA using endonuclease-mediated ssDNA conjugation and photo-crosslinking of protein G. A pG-HUH tag fusion protein is reacted with ssDNA containing an HUH recognition sequence by simply mixing. Subsequently, an antibody is added to the mixture and exposed to UV light to enable covalent photo-crosslinking to the pBPA unnatural amino acids present in the protein G domains. The antibody-DNA conjugate can be used for several applications in nanotechnology or diagnostics.

## Results and discussion

### Construct design & protein expression

Labeling antibodies with photo-crosslinkable protein G usually yields a mixture of mono- and bi-labelled species.^13,28^ We recently reported the construction of a protein G dimer consisting of two protein G domains connected by a peptide linker that spans the distance between the two heavy chains of an antibody. This configuration enabled efficient conjugation of split luciferase domains in a precise 1:1 ratio.^29^ Here, we use this protein G dimer to site-specifically label antibodies with DNA, improving on previous methods that yielded a mixture of antibodies containing one or two DNA strands.^13,14^ The protein G dimer was fused to one of three structurally characterized Reps used in previous studies: DCV (*muscovy duck circovirus*), PCV2 (*porcine circovirus 2*) and WDV (*wheat dwarf virus*). Because introducing unnatural amino acids can lead to premature termination of protein synthesis, purification tags were introduced at the N- and C- termini of the fusion proteins. The pG-Rep fusion proteins were expressed in *E. coli* and purified by Ni^2+^ affinity and Strep-Tactin® affinity chromatography. Despite the incorporation of two unnatural amino acids, the purified fusion proteins were obtained in high yields: 24 mg/L for pG-DCV, 18 mg/L for pG-PCV2, and 4.8 mg/mL for pG-WDV (Figure S2). Protein identity and correct incorporation of pBPA were confirmed using QToF mass spectrometry (Figure S3).

### HUH-DNA conjugation characterization

Rep proteins recognise specific ssDNA sequences and cleave them by forming a covalent phosphate ester bond with the 5’ end of the resulting 3’ cleavage fragment (Figure 2A). To evaluate the efficiency and specificity of DNA conjugation, we started with a consensus ssDNA sequence (Seq1) previously reported to be efficiently cleaved by all three Rep proteinsin the presence of Mn^2+^ (Table S1). Sequence specificity was tested by introducing single or multiple mismatches at various positions within Seq1 (Table S1, Table S2). Previous studies using DCV showed increased affinity and a 4-fold faster enzymatic activity when using Mn^2+^ as cofactor instead of Mg^2+. 24^ Since most prior studies used Mn^2+^ as a cofactor, we compared Mg^2+^ and Mn^2+^ to determine whether the presence of Mg^2+^ might enhance sequence specificity. In these experiments, 4 µM pG-Rep was incubated with a 10-fold molar excess of ssDNA (either Seq1 or mismatched variant), in the presence of 1 mM Mg^2+^ or 1 mM Mn^2+^ (Figure 2A). Reactions were incubated overnight and analysed the following day by SDS-PAGE, stained with SYBR™ Gold to visualize DNA and Coomassie blue to visualize proteins. In the presence of Mg^2+^, complete conjugation of Seq1 was observed for pG-DCV (Figure 2B) and pG-PCV2 (Figure 2C), while pG-WDW showed only partial conjugation (Figure 2D). Conjugation was highly specific in the presence of Mg^2+^, as no significant conjugation was observed for sequences containing one or more mismatches. In contrast, Mn^2+^ showed decreased sequence specificity. All three domains showed essentially complete conjugation with Seq1 with Mn^2+^ as cofactor, but also substantial conjugation to mismatched sequences. Based on these results, we proceeded with Mg^2+^ as the cofactor in subsequent experiments.

**Figure 2.**
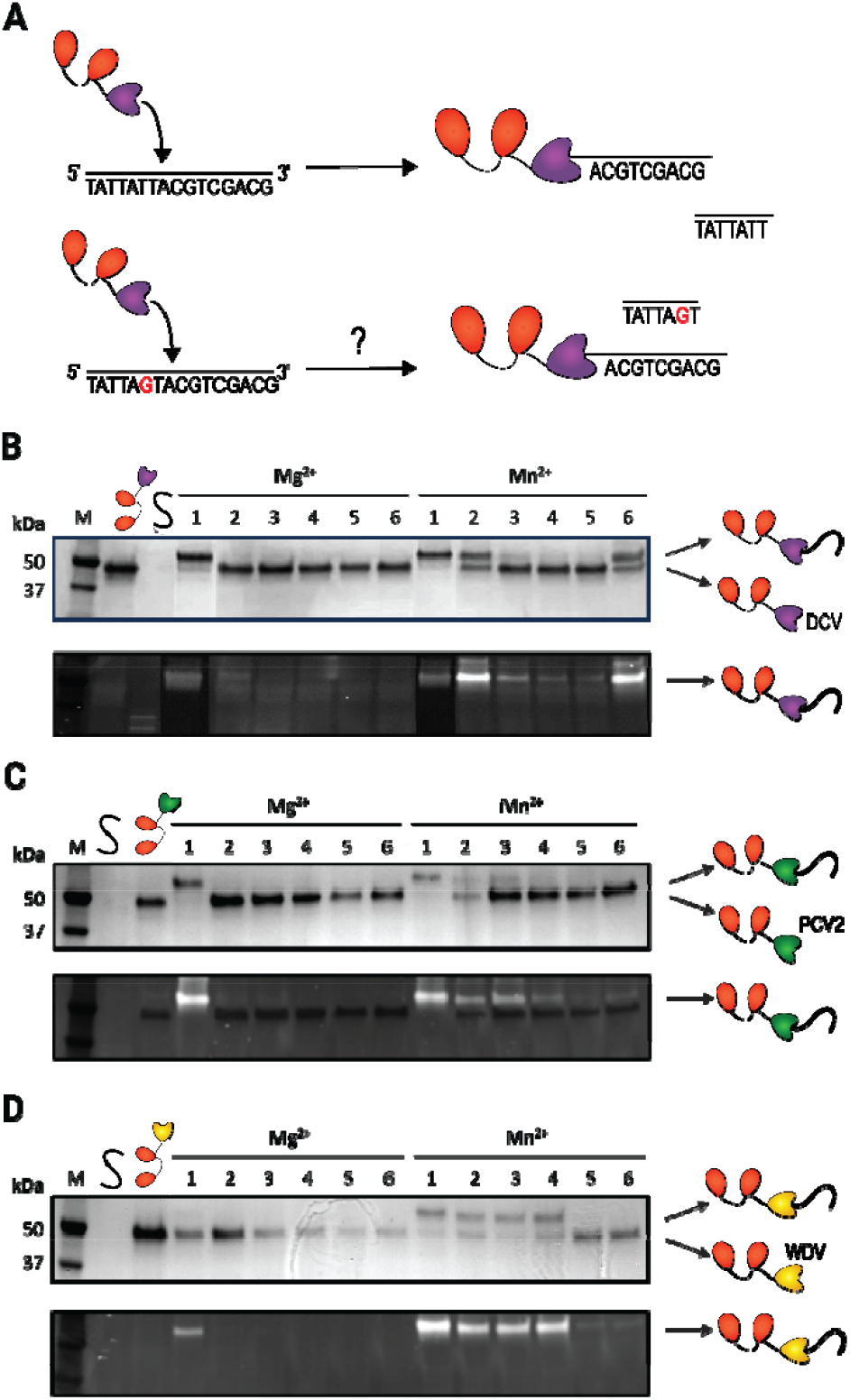
Characterization of DNA recognition by Rep proteins. **A)** Graphical representation of the experiment. The fusion proteins pG-HUH were reacted with 10 equivalents of ssDNA. **B-D)** Experiments investigating the sequence specificity of **B)** pG-DCV, **C)** pG-PCV2, **D)** pG-WDV. The top panel is the Coomassie-stained SDS-PAGE gel to visualise proteins, whereas the lower panel is the same gel stained with SYBR™ Gold to visualize DNA. The experiment showcases 6 different ssDNA in the presence of either Mg^2+^ or Mn^2+^. Sequence 1 (noted 1 in the figure) is predicted to be cleaved by all endonucleases. Sequences 2-6 contain point mutations in various positions of Seq1.

### pH dependence and phosphotyrosine bond stability

Phosphotyrosine bonds, such as formed here between endonucleases and DNA, are known to be reversible and may be susceptible to hydrolysis, especially under basic or acidic conditions. To investigate the pH sensitivity of the conjugation reaction, pG-HUH fusion proteins were incubated with a 10-fold molar excess of Seq1 ssDNA at pH 6.0, 7.5 and 9.0 and samples were taken after 1 hour, 1 day, and 2 days. Figure 3 shows that the conjugation reaction is complete within 1 hour for both pG-DCV and pG-PCV2 at all tested pH values. Moreover, no hydrolysis was detected after 2 days of incubation. In contrast, conjugation of pG-WDV (Figure 3D) was clearly pH dependent, with little conjugation observed at pH 6. Consistent with the results in Figure 2, conjugation of Seq1 with pG-WDV was also much less efficient at pH 7.5 and 9, requiring 1-2 days to achieve ∼80% conversion.

**Figure 3.**
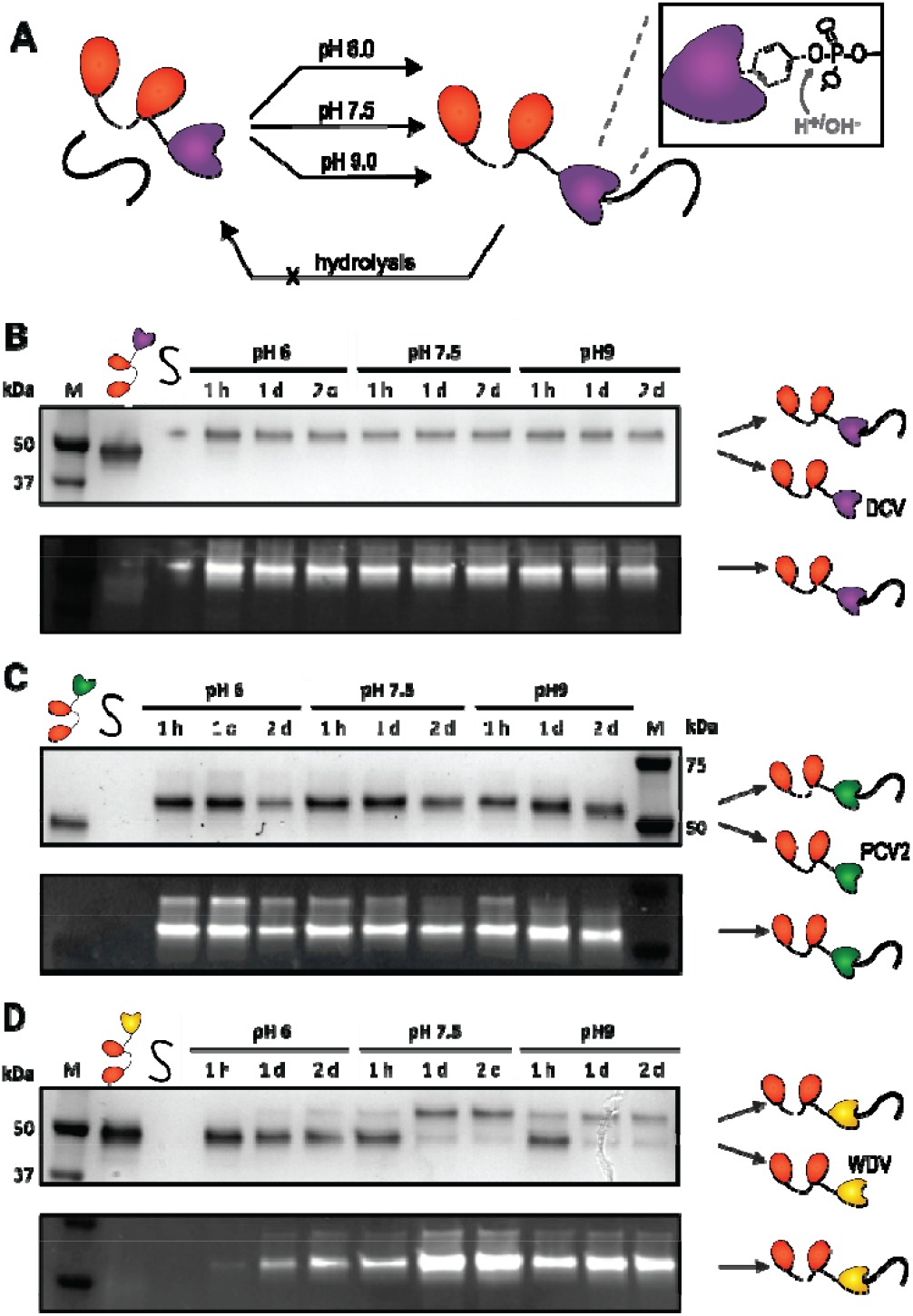
Robustness of the bioconjugate formation and stability at various pH. **A)** Graphical representation of the experiment. The pG-HUH was reacted with 10 equivalents of ssDNA in buffers of pH 6, 7.5 and 9. **B-D)** The pG-HUH fusion proteins: **B)** pG-DCV, **C)** pG-PCV2, **D)** pG-WDV, were reacted with ssDNA at three different pH conditions and studied over the course of 2 days (2d). The top panel is the Coomassie-stained SDS-PAGE gel to visualize proteins, whereas the lower panel is the same gel stained with SYBR™ Gold to visualize DNA.

Although no hydrolysis was detected after 2 days incubation, these experiments were performed in the presence of a 10-fold molar excess of ssDNA Seq1. Therefore, the results do not exclude the possibility that the phosphotyrosine undergoes hydrolysis, followed by reaction of the Rep protein with another ssDNA strand. To test this possibility, we performed an assay in which pG-DCV was first reacted with a 3-fold molar excess of a short ssDNA oligo for 1 hour at pH 7.5. Subsequently, a second, longer ssDNA oligo containing the same recognition sequence was added in 3-, 5- or 10-fold molar excess and incubated overnight. The use of DNA of different lengths enabled straightforward detection of strand exchange by SDS-PAGE. A similar assay was performed with pG-PCV2 and pG-WDV (Figure S4A-C), along with a complementary experiment in which pG-HUH was first reacted with the longer DNA strands and then challenged overnight with the shorter DNA strand (Figure S4D-G). In both setups, all Reps showed exchange of DNA strands, irrespective of the length of the competing DNA. In addition, increasing the amount of competing strand increased the amount of exchange.

These strand exchange experiments suggest that formation of the tyrosine-phosphate bond is reversible, most likely through hydrolysis of the initially formed DNA conjugate, followed by subsequent cleavage and re-conjugation of another ssDNA containing the recognition sequence. To more stringently assess the stability of the HUH-DNA conjugates for the DCV protein, 4 µM of pG-DCV was reacted with a 10-fold molar excess of ssDNA containing the recognition sequence. The resulting pG-DCV-DNA conjugate was purified using StrepTactin® affinity chromatography to remove unreacted DNA (Figure 4C). The purified conjugate was incubated at pH 7.5 (100 mM Tris-HCl) at room temperature, and samples were analyzed by SDS-PAGE over a period of 8 days (Figure 4D). Under these conditions, the pG-DCV-DNA remained stable, with no detectable band corresponding to free pG-DCV throughout the 8 days experiment. We therefore conclude that although strand exchange experiments suggest that the tyrosine-phosphate bond can be hydrolyzed, the rate of hydrolysis is low, and the phosphotyrosine bond can be considered stable under these conditions.

**Figure 4.**
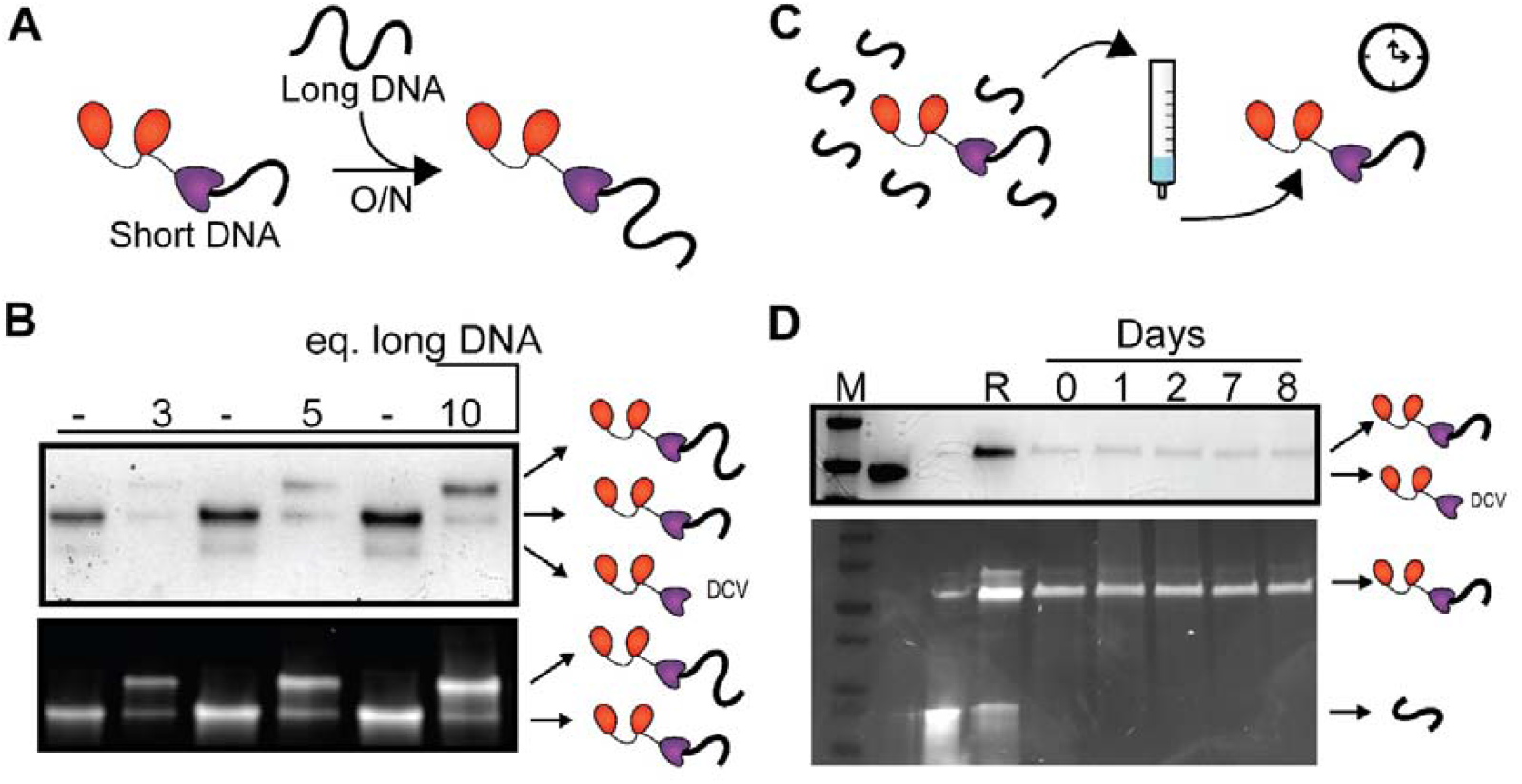
Reversibility of phosphotyrosine bond formation. **A)** Graphical representation of the competition assay between ssDNA containing the recognition sequence. Shorter ssDNA is reacted with pG-DCV and subsequently, longer ssDNA is added for overnight incubation. **B)** SDS-PAGE displaying pG-DCV pre-reacted with 3 equivalents of short DNA, followed by the addition of long DNA in 3, 5, 10 molar equivalents. **C)** Graphical representation of the experiment. pG-DCV is reacted with the ssDNA and subsequently purified on the StrepTactin® resin. The purified construct is kept at room temperature over the course of 8 days. **D)** SDS-PAGE displaying the respective samples R: reaction mix, 0: the purified conjugate, 1-8: number of days after the purification. The top panels are the Coomassie-stained SDS-PAGE gel to visualize proteins, whereas the lower panels are the same gel stained with SYBR™ Gold to visualize DNA.

### Synthesis of DNA-antibody conjugates for proximity extension assays

Having established the stability of the pG-DCV-DNA conjugates, we next used them to generate antibody-DNA conjugates via photocrosslinking. As an initial application, we chose to develop antibody-DNA conjugates for the detection of 2 clinically relevant proteins, TNFα and interleukin-6 (IL-6), using proximity extension assays (PEA). For IL-6 detection, we used two monoclonal antibodies R508 and mhK23, that bind to different epitopes and form an established sandwich pair. In contrast, because TNFα exists as a homotrimer, a single antibody (Adalimumab) was sufficient for detection of this protein. We used the same pair of oligos (S1 and S2, Table S3), each containing a DCV recognition sequence at the 5’ end and a complementary six nucleotide sequence at the 3’ end, which was adopted from a previously reported design.^2^ This complementary sequence is too short to hybridize when the antibody-DNA conjugates are free in solution. However, when antibodies carrying S1 and S2 bind their target protein, they are brought into close proximity, enabling hybridization. In PEA, this hybridized complex is first extended by T4 DNA polymerase, generating a dsDNA product, which is then quantified using qPCR. To prepare the conjugates, 7.5 µM pG-DCV was reacted with various amounts of S1 and S2 ssDNA (Figure 5B). Complete conjugation was achieved at a 1:1 molar ratio, eliminating the need for purification to avoid background from excess DNA. The resulting pG-DCV–S1 and pG-DCV–S2 products from the 1:1 reaction were directly used in subsequent antibody conjugations. For these reactions, 1 µM of R508 and mhK23 antibodies was mixed with 1- to 4-fold molar excess of the pG-DCV-S1 or pG-DCV-S2, respectively, and irradiated with 365 nm UV light for 30 min. As shown in Figures 5C-D, both reactions resulted in the efficient formation of single antibody-DNA conjugates in 60-80% yield. Comparable results were obtained when S1 and S2 were conjugated to Adalimumab (Figure S5). Based on these results, a 2:1 molar ratio of pG-DCV-DNA to antibody was used in the PEA experiments.

**Figure 5.**
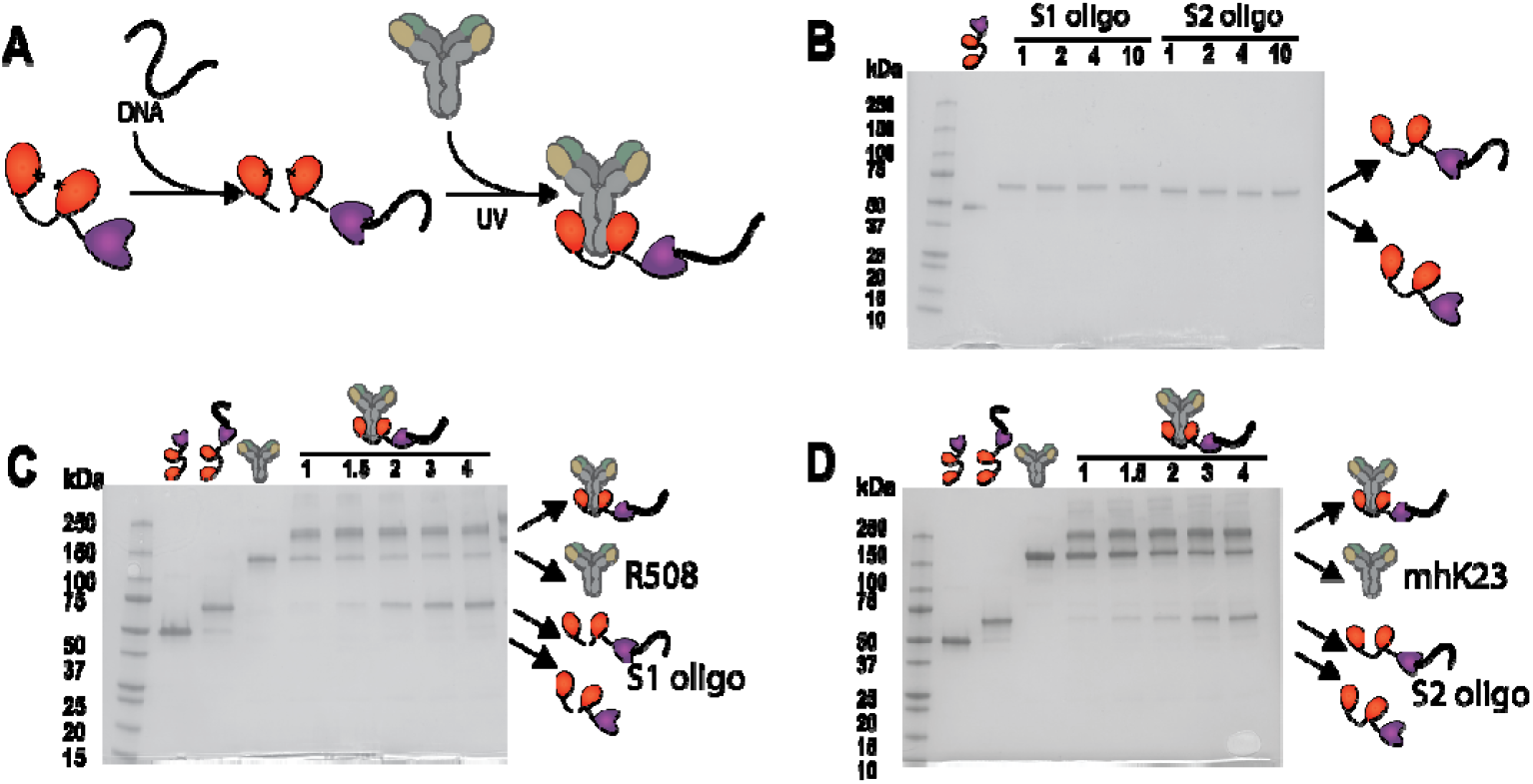
Photocrosslinking of pG-DCV to the anti-IL-6 antibodies R508 and mhK23. A) Graphical representation of the experiment. pG-DCV was reacted with ssDNA and subsequently photo-crosslinked to the antibody. B) SDS-PAGE showing the reaction of pG-DCV with 1, 2, 4 or 10 molar equivalents of S1 or S2 DNA. C-D) SDS-PAGE gel monitoring the photoconjugation of R508 and mhK23 antibodies using 1- to 4-fold molar excess of pG-DCV-DNA compared to the antibody.

To perform PEA, 10 pM of the anti-IL-6 or Adalimumab conjugates were incubated with increasing concentrations of IL-6 or TNFαfor 30 min at 37 °C. Subsequently, T4 polymerase was added and incubated for 90 min at 37 °C, after which the reaction mixtures were analyzed by qPCR (Figure 6A). Specific detection of IL-6 and TNFα was observed using the respective antibody-DNA conjugates, whereas cycle threshold (Cq) values for non-binding antibody-DNA conjugates were similar to the background observed in the absence of target analyte. Titration experiments showed that concentrations as low as 24 pM IL-6 and 49 pM TNFα could be reliably detected above background (Figure S6). These results confirm that the antibody-DNA conjugates are functional and that DNA-conjugation does not impede antibody binding. The relatively modest change in Cq value observed in these assays may be further improved by applying a fluorophore-quencher DNA probe specific to the amplified target, instead of using a generic dsDNA-intercalating dye.

**Figure 6.**
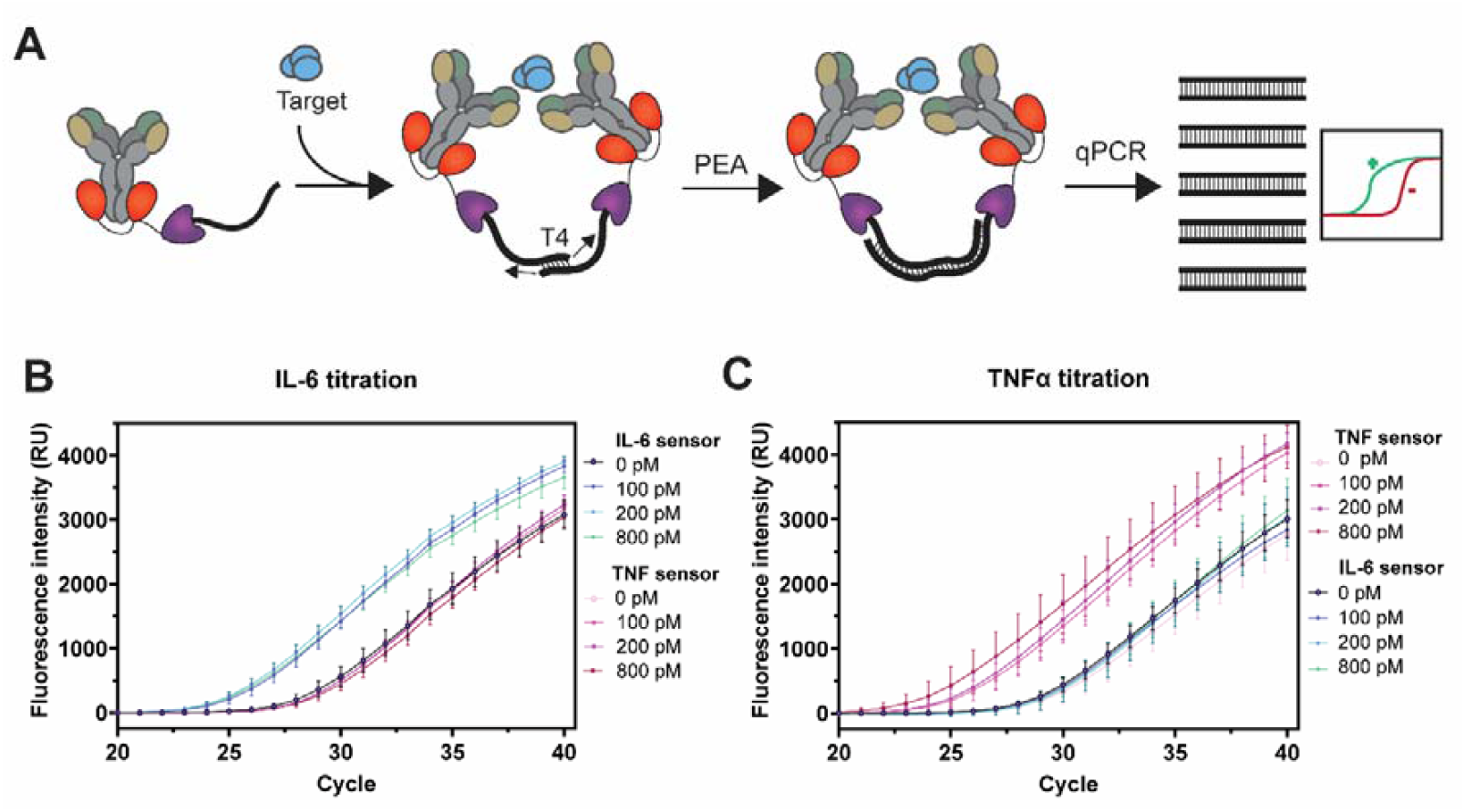
Use of antibody-DNA constructs for PEA assays detecting TNFα and IL-6. A) Graphical representation of the experiment. Two antibodies binding to the same molecule allow the hybridization of complementary DNA-strands with free 3’-OH groups, which allows for subsequent DNA extension by T4 Polymerase. The amplified DNA fragments can be quantified using qPCR. B-C) PEA dose-response curves for Interleukin-6 (B) and TNFα (C). qPCR experiments were performed in triplicate.

## Conclusions and discussion

This work introduced a new site-specific method for covalently attaching single-stranded DNA to the Fc domain of antibodies using a fusion protein composed of a protein G dimer and an HUH endonuclease. Among three HUH variants tested (DCV, PCV2, WDV), the DCV domain provided efficient, nearly complete DNA conjugation at a 1:1 protein-to-DNA ratio without requiring excess oligonucleotides. DNA attachment showed higher sequence specificity with Mg^2+^ compared to Mn^2+^ and remained stable for at least eight days at room temperature. The resulting antibody–DNA conjugates, after photo-crosslinking, were functional, as proven in proximity extension assays for TNFα and IL-6 detection.

The simple and fast conjugation approach, requiring only unmodified oligos and a readily expressed pG-HUH fusion protein, is broadly applicable beyond PEA. It is particularly attractive for applications that involve the development and screening of a large number of antibody-DNA conjugates, such as immuno-PCR and multiplex DNA-PAINT. The site-specific conjugation to the Fc-part ensures undisturbed antigen binding, which together with the precise 1:1 labelling stoichiometry is critical for quantitative imaging applications. Mono-labelled antibodies are also advantageous for antibody functionalization of DNA Origami structures, where they mitigate aggregation by preventing multivalent attachment of an antibody to multiple DNA structures.

Although photo-cross-linkable protein G allows conjugation to several classes of mammalian IgGs and all human IgG isotypes, protein G-mediated conjugation is incompatible with several important antibody formats including mouse IgG1, avian IgY, and scFv and Fab antibody fragments. In these cases, a truncated variant of protein M (pM440) could be used instead of protein G. pM440 binds to the light chain of a broad range of antibodies with sub-nanomolar affinity. We recently reported the successful substitution of protein G-mediated photo-conjugation by pM440 in bioluminescent sandwich immunoassays based on split luciferase complementation.^30^ In that work, pM440 antibody complex formation was found to be essentially irreversible, eliminating the need for photo-crosslinking.

## Methods

### Molecular Cloning of pG-HUH constructs

All gblocks were purchased at IDT. The pET28 plasmid encoding the pG-DCV was constructed via restriction/ligation. DCV gblock was cloned C-terminally of pG originating from pet28a pG-Dimer-LgBiT our group recently described ^31^ using Nco1, Not1 and Bbs1. Thereby, a single AgeI restriction site was introduced. The other plasmids pET28 pG-PCV2, and pG-WDV were cloned by performing restriction with AgeI, and BamHI on the gBlock coding for PCV2 or WDV, and pG-DCV including rSAP (Shrimp Alkaline Phosphatase) to prevent self-ligation. The restriction reaction of the plasmid was analyzed with agarose gel electrophoresis. Gel extraction (QIAquick Gel Extraction Kit, #28706) was performed on the backbone, while PCR cleanup (QIAquick PCR Purification Kit, #28106) was performed on the gBlocks. The backbone and insert were ligated using T4-ligase for 1.5 hours at 16 °C. The ligation product was transformed into heat competent *Escherichia coli* Top10 cells. The plasmids were extracted from the bacteria with MiniPrep (QIAprep Spin Miniprep Kit, #12123). The successful cloning was confirmed with Sanger Sequencing (BaseClear).

### Expression of pG-HUH proteins

The plasmids, containing sequences of pG-DCV, pG-PCV2, or pG-WDV were co-transformed with pEVOL-pBpF in *Escherichia coli* BL21 (DE3). The pEVOL vector encodes the tRNA and tRNA synthetase, which incorporates the pBPA at the amber stop codon.^32^ The cells were cultured in 500 mL LB medium (5 g tryptone, 5 g NaCl, and 2.5 g Yeast extract) containing 50 µg/mL kanamycin and 25 µg/mL chloramphenicol. Cells were grown at 37 °C and 200 rpm until an OD600 of 0.6-0.9 and induced with 0.8 mM IPTG, 0.02% arabinose, and 1 mM pBPA (Bachem, #104504-45-2). The expression was conducted overnight at 18 °C and 160 rpm shaking. The cells were harvested by centrifugation at 8000 g for 5 minutes. Lysis was carried out with the Bugbuster reagent (Novagen, #70584) and 5 µL benzonase (Novagen, #70746-3).

### Protein purification

All proteins were purified using the hexahistidine and StrepTag II tags. The Ni-NTA column was loaded with Ni-NTA agarose (Qiagen, #30230). The column was equilibrated with Buffer A (50 mM Tris-Cl pH 8, 500 mM NaCl, and 10 mM imidazole). The lysate was applied to the column, the column was washed with Buffer A and the protein was eluted with elution buffer (50 mM Tris-HCl pH 8, 500 mM NaCl, and 350 mM imidazole). The strep column was loaded with Strep-Tactin® XT 4Flow® high capacity resin (IBA lifesciences, #2-5030-025). The column was equilibrated with equilibration buffer (100 mM Tris-Cl pH 8, 150 mM NaCl, and 1 mM EDTA). The elution of the Ni-NTA column was added to the StrepTactin column. The column was washed with the equilibration buffer and the proteins were eluted with the elution buffer (100 mM Tris-Cl pH 8, 150 mM NaCl, 1 mM EDTA, and 50 mM D-biotin). Purification, correct expression of the protein, and incorporation of pBPA were validated with SDS-PAGE and Q-ToF.

### Q-ToF

Purified proteins were buffered exchanged to 0.1% formic acid in Milli-Q using PD SpintrapTM G-25 column (Cytiva, #28918004). The Q-ToF measurement was performed on the Q-ToF LC-MS (WatersMassLynx v4.1). MagTran v1.03. was used to deconvolute the data.

### HUH endonuclease ssDNA conjugation

pG-DCV, pG-PCV2, and pG-WDV were conjugated to varied strands of ssDNA (IDT, Table S1-3). Typically, conjugation was performed with 4 µM HUH endonuclease and 10-fold excess of ssDNA (unless stated otherwise) in conjugation buffer (50 mM HEPES pH 7.5, 50 mM NaCl, 1 mM MgCl_2_) and incubated for one hour at room temperature or overnight. In the experiments comparing the influence of Mg^2+^ and Mn^2+^, 1 mM of metal ion was used. In the experiment comparing pH, 50 mM of Bis-Tris pH 6.0 or 50 mM of Tris pH 9.0 were used instead of HEPES buffer. For conjugate purification on the Strep-Tactin column, the same method as for protein purification was applied.

### Photocrosslinking

4 µM of pG-DCV was reacted with 4 µM of ssDNA in conjugation buffer and incubated overnight. Subsequently, the reaction was diluted to 2 µM and 1 µM of antibody was added and the mixture was incubated on ice for 30 minutes. The photocrosslinking reaction was performed for 30 minutes at 50% intensity with a continuous wave using a 365 nm UV-lamp (Thorlabs, #M365LP1) and T-Cube LED Driver (Thorlabs, #LEDD1B). The photocrosslinking reaction was typically analyzed with a non-reducing SDS-PAGE.

### SDS-PAGE

Samples were mixed with 2x SDS-PAGE Laemmli buffer (125 mM Tris-Cl pH 6.8, 20% (v/v) glycerol, 5% SDS, 0.02% bromophenol blue), either in the presence of 50 mM DTT (reducing for pG-Rep-DNA conjugation) or absence (not reducing for photocrosslinking experiments) and boiled for 5-10 minutes at 95 °C. The samples (or protein ladder (Bio-Rad, #1610373)) were applied to the precast gel (4-20% Mini-PROTEAN® TGX, Bio-Rad, #4561096 or #4561093) and run in running buffer (25 mM Tris, 192 mM glycine, 0.1% (v/v) SDS, Bio-Rad, #1610772). The gel was washed with Milli-Q for 30 minutes, stained with 5 µL 1000x SYBR gold (ThermoFisher, #S11494) for 20 minutes, and washed with Milli-Q for an additional 5-20 minutes. The gel was imaged with Cytiva Amersham Image Quant TM 800 at 470 nm. Afterwards, the gel was stained with Coomassie (Bio-safe Coomassie G-250 stain, Bio-Rad, #1610786) for 30 minutes and washed with Milli-Q. The same imager, was used to visualize the Coomassie staining.

### Proximity extension assay (PEA)

Antibody-DNA conjugates were diluted to 1 nM in buffer (1x PBS, 0.1 % BSA). The PEA reaction mixture consisted of 10 pM of antibody-DNA conjugates, 40 µM dNTPs, 0.016 mg/mL salmon sperm DNA and 1x NEB Buffer™ r2.1 (NEB, #B6002S). The target was added and the mixtures were incubated for 30 min at 37 °C. 0.5 µL of T4 polymerase (NEB, #MO203S) was added and incubated for an additional 90 min at 37 °C. The polymerase was inactivated for 10 min at 80 °C. 1 µL of the reaction was transferred to the qPCR mixture consisting of 1x Taq polymerase Master Mix (NEB, #M0270L), 1x EvaGreen® Plus dye (Biotium, #31077-T) and 200 nM of primers (Table S3). The qPCR cycles consisted of an initial denaturation step of 5 min at 95 °C, followed by 40 cycles: 95 °C for 1 min, 52 °C for 1 min.

## Acknowledgments

The authors would like to thank Adam Smiley and Matthew Pawlak from the Gordon group for insightful discussions. We would like to thank the members of the Merkx group for fruitful discussions and feedback.

