## supporting information for "Homogeneous antibody-DNA-conjugates using unmodified oligonucleotides and photo-crosslinkable protein G-HUH endonuclease fusion proteins"

| **Supplementary Figures:** | | Page |
|---|---|---|
| Table S1 | Predicted cleavage based on HUH-seq screening | S3 |
| Table S2 | DNA sequences used in a specificity assay | S3 |
| Table S3 | DNA used for proximity extension assay | S4 |
| Figure S1 | Protein sequences | S4 |
| Figure S2 | Purification of used fusion proteins pG-Rep | S5 |
| Figure S3 | Qtof analysis of purified proteins | S5 |
| Figure S4 | The competition assay between DNA containing the recognition sequence | S6 |
| Figure S5 | SDS-PAGE displaying the required excess of pG-DCV-DNA over Adalimumab for PEA | S7 |
| Figure S6 | Figure S6 PEA dose-response curves for Interleukin-6 and TNFα | S7 |

Supplementary references S7

**Supplementary Figure and Tables:**

**Table S1 Predicted cleavage based on HUH-seq screening using $Mn^{2+}$ as a cofactor.** [1] The summary is based on the HUH endonuclease and ssDNA used in the specificity experiment. The provided log2FC values denote the predicted cleavage. With the log2FC value less than -3, HUH endonuclease is predicted to recognize the sequence and perform cleavage. If the log2FC value exceeds -0.3, no cleavage is predicted. With intermediate value, cleavage can occur to various degree. Retrieved from reference 1.

| **Recognition sequence** | **DCV** | **WDV** | **PCV2** |
|---|---|---|---|
| 1. TAT TAT TAC | -3.89 | -4.07 | -4.18 |
| 2. TAT **G**AT TAC | -0.46 | -0.55 | -2.16 |
| 3. TA**A G**AT TAC | -0.16 | -3.74 | -1.12 |
| 4. TA**A GC**T TAC | -0.13 | -2.32 | -0.65 |
| 5. T**GA GC**T TAC | 0.02 | -0.14 | -0.13 |
| 6. TAT TAT T**GT** | Not studied | Not studied | Not studied |

**Table S2 DNA sequences used in a specificity assay.** The asterisk denotes the cleavage site. As the initial experiment (Figure 2) did not show full conversion of WDV with recognition sequence (Seq1), another ssDNA was tested for WDV (Seq9-10). The mismatched positions are bolded.

| **ssDNA strand** | **Sequence** |
|---|---|
| 1. recognition sequence | TAT TAT T*AC CGC TT CGT CTG CTT CAT CGC TGT TA |
| 2. 1 mismatch | TAT **G**AT T*AC TT CGT CTG CTT CAT CGC TGT TA |
| 3. 2 mismatches, -4, and -5 positions | TA**A G**AT T*AC TT CGT CTG CTT CAT CGC TGT TA |
| 4. 3 mismatches | TA**A GC**T T*AC TT CGT CTG CTT CAT CGC TGT TA |
| 5. 4 mismatches | T**GA GC**T T*AC TT CGT CTG CTT CAT CGC TGT TA |
| 6. 2 mismatches, +1, and +2 | TAT TAT T***GT** TT CGT CTG CTT CAT CGC TGT TA |
| 7. reversibility short ssDNA, DCV, and PCV2 | TAT TAT T*AC CTT TAC CAA CTT CGC ATA CTA CTA C |
| 8. reversibility long ssDNA, DCV, and PCV2 | TAT TAT T*AC CTT CAA CTT CGC ATA CTA CTA CTA CTA CTA CTA CTA CTA CTA CTA CTA C |
| 9. reversibility short ssDNA, WDV | TAA TTT T*AC CTC TAC CAA CTT CGC CCA CTA CTA C |
| 10. reversibility long ssDNA, WDV | TAA TTT T*AC CTC TAC CAA CTT CGC CCA CTA CTA CTA CTA CTA CTA CTA CTA CTA CTA C |

**Table S3 DNA used for proximity extension assay.**

| ssDNA | Sequence |
|---|---|
| 1. S1 | TATTATTACTTTTGTTGAGGTAACCAACTATTTGTT ACTGTTGCGTACGTCGCCGTCCAG |
| 2. S2 | TATTATTACTTTTCCTCACCATGTCTGAGGTACTC CTTAAAGGTCGAGCTGGAC |
| 3. Forward qPCR primer | TTGTTGAGGTAACCAACTATTTGTTACTGTT |
| 4. Reverse qPCR primer | TTCCTCACCATGTCTGAGGTACTCCTTAAAGG |

**A**

MGWSHPQFEKGGSMTFKLIINGKTLKGEITIEAVDA*EAEKIFKQYANDYGIDGEWTYDDATK
TFTVTEEFTGGSGGSGGSGGSGGSGGSGEFAEAAAKEAAAKEAAAKEAAAKEAAAKEAAA
KAEFGGSGGSGGSGGSGGSGGSGGTMTFKLIINGKTLKGEITIEAVDA*EAEKIFKQYANDY
GIDGEWTYDDATKTFTVTELTGGSGGSGGSGGSGGSGGSGEFAEAAAKEAAAKEAAAKEA
AAKEAAAKEAAAKAEFGGSGGSGGSGGSGGSGGSGGTGMAKSGNYSYKRWVFTINNPTF
EDYVHVLEFCTLDNCKFAIVGEEKGANGTPHLQGFLNLRSNARAAALEESLGGRAWLSRAR
GSDEDNEEYCAKESTYLRVGEPVSKGRSSGSGGSHHHHHH

**B**

MGWSHPQFEKGGSMTFKLIINGKTLKGEITIEAVDA*EAEKIFKQYANDYGIDGEWTYDDATK
TFTVTEEFTGGSGGSGGSGGSGGSGGSGEFAEAAAKEAAAKEAAAKEAAAKEAAAKEAAA
KAEFGGSGGSGGSGGSGGSGGSGGTMTFKLIINGKTLKGEITIEAVDA*EAEKIFKQYANDY
GIDGEWTYDDATKTFTVTELTGGSGGSGGSGGSGGSGGSGEFAEAAAKEAAAKEAAAKEA
AAKEAAAKEAAAKAEFGGSGGSGGSGGSGGSGGSGGTGMASSSTPRFRVYSKYLFLTYP
QCTLEPQYALDSLRTLLNKYEPLYIAAVRELHEDGSPHLHVLVQNKLRASITNPNALNLRMDT
SPFSIFHPNIQAAKDCNQVRDYITKEVDSDVNTAEWGTFVAVSTPGRKDRDADGSGGSHHH
HHH

**C**

MGWSHPQFEKGGSMTFKLIINGKTLKGEITIEAVDA*EAEKIFKQYANDYGIDGEWTYDDATK
TFTVTEEFTGGSGGSGGSGGSGGSGGSGEFAEAAAKEAAAKEAAAKEAAAKEAAAKEAAA
KAEFGGSGGSGGSGGSGGSGGSGGTMTFKLIINGKTLKGEITIEAVDA*EAEKIFKQYANDY
GIDGEWTYDDATKTFTVTELTGGSGGSGGSGGSGGSGGSGEFAEAAAKEAAAKEAAAKEA
AAKEAAAKEAAAKAEFGGSGGSGGSGGSGGSGGSGGTGPSKKNGRSGPQPHKRWVFTL
NNPSEDERKKIRDLPISLFDYFIVGEEGNEEGRTPHLQGFANFVKKQTFNKVKWYLGARCHI
EKAKGTDQQNKEYCSKEGNLLMECGAPRSQGQRGSGGSHHHHHH

**Figure S1 Protein sequences of protein G dimer fused to A) DCV B) WDV C) PCV2**. The asterisk represents the Amber stop codon for the introduction of an unnatural amino acid. The highlighted sequences are StrepTag® II, protein G dimer, HUH tag and 6xHis tag.

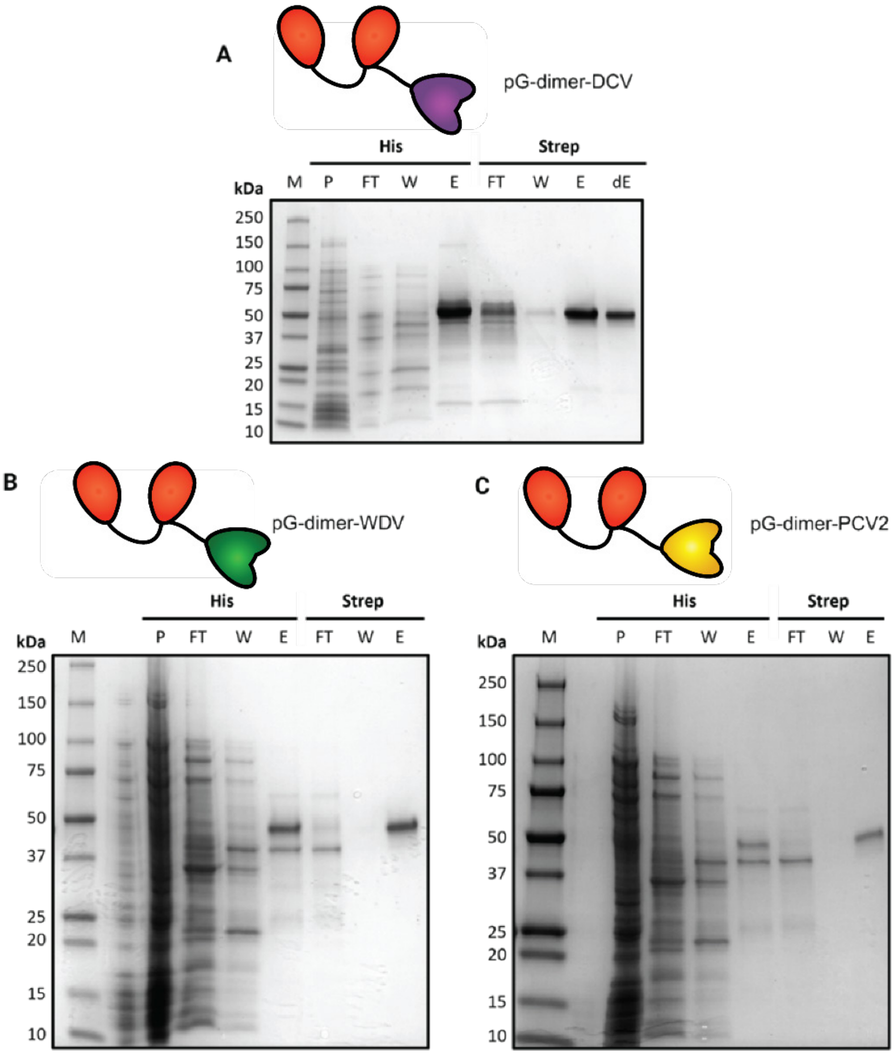


**Figure S2 Protein purification of protein G dimer fused to A) DCV, B) WDV, C) PCV2.** The purification was conducted in two steps: first to capture the Hexahistidine tag (His) and second to capture the strep-tag (Strep). The lanes are as follows: M: Protein marker, P: Pellet, FT: Flow through, W: Wash, E: Elution.

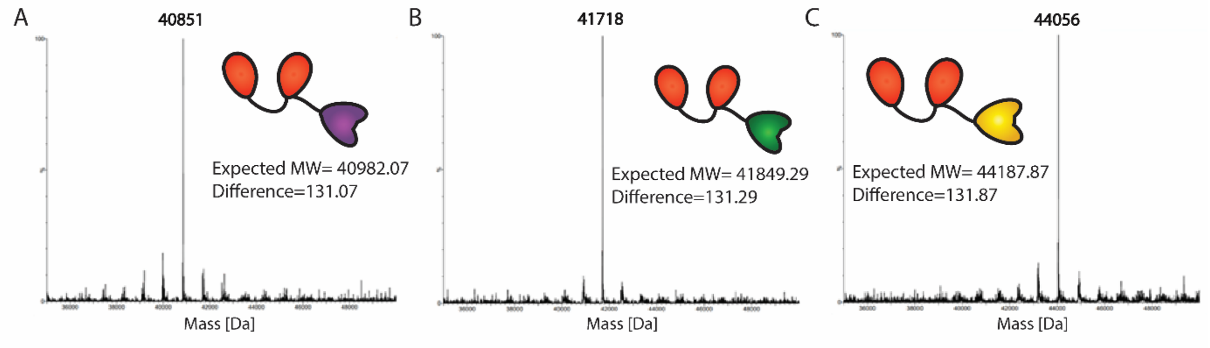


**Figure S3 QToF analysis of protein G dimer fused to A) DCV, B) PCV2, C) WDV.** Fusion proteins were buffer exchanged to 0.1% formic acid in Milli-Q and analyzed by LC-MS. The observed mass profiles matched the expected values, accounting for the typical cleavage of N-terminal (formyl)methionine in *E. coli* (131 Da).

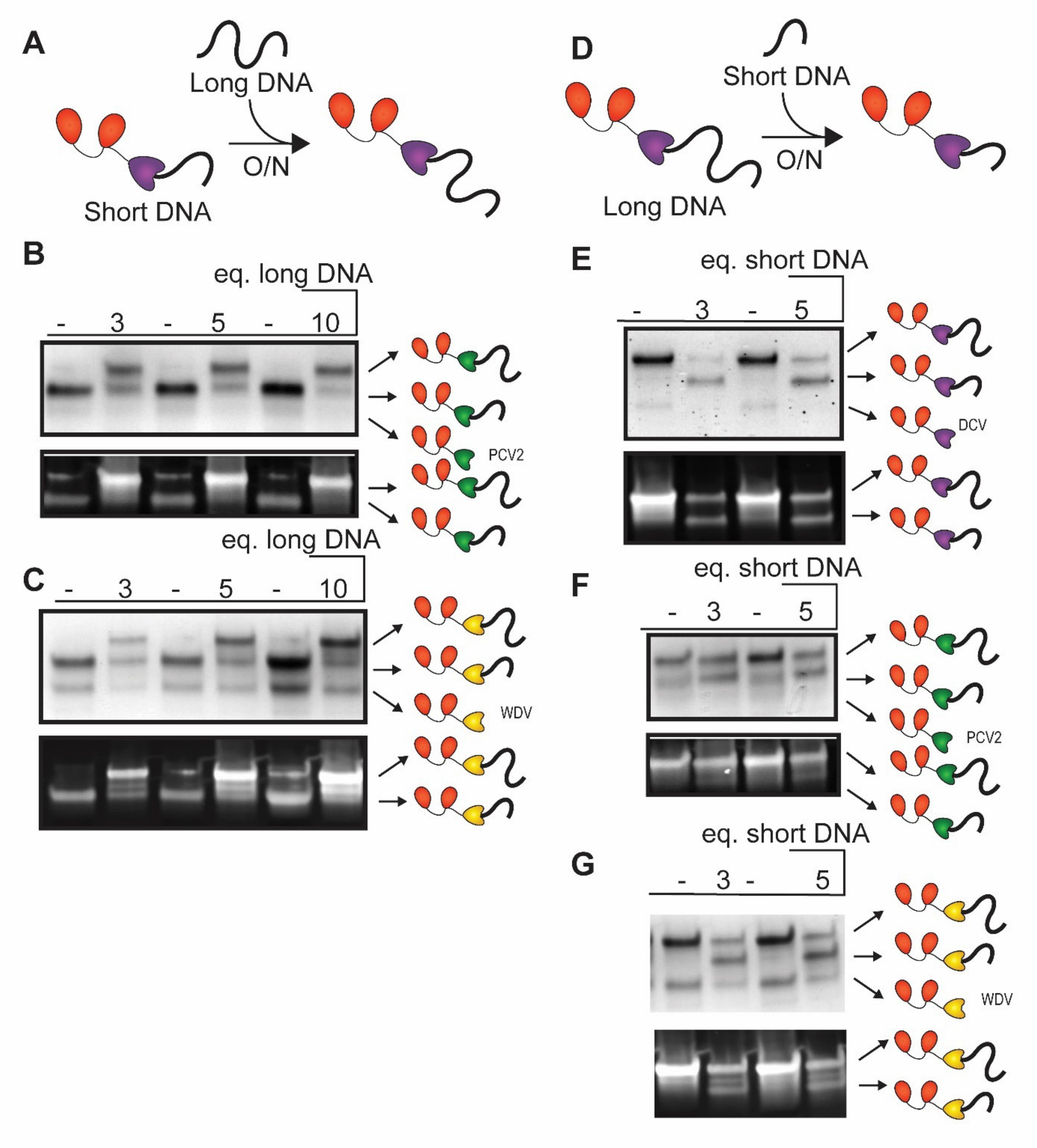


**Figure S4 DNA exchange assay.** A) 4 µM pG-Rep construct was reacted with 3 molar equivalents of ssDNA oligo for 1 h. Next, 3, 5 or 10 molar equivalent excesses of longer (A-C) or shorter (D-G) ssDNA containing the recognition sequence were added and allowed to compete with the initial oligo overnight. The following fusion proteins were studies: pG-DCV (Figure S4E), PCV2 (Figure S4B, S4F), and WDV (Figure S4C, S4G). After the overnight reaction, the intensity of the initial pre-conjugated band decreases and another band appears, indicating the formation of the conjugate with other ssDNA. ssDNA sequences are listed in Table S2. The top panel is the Coomassie stained SDS-PAGE gel to visualize proteins, whereas lower panel is the same gel stained with SYBR™ Gold to visualize DNA.

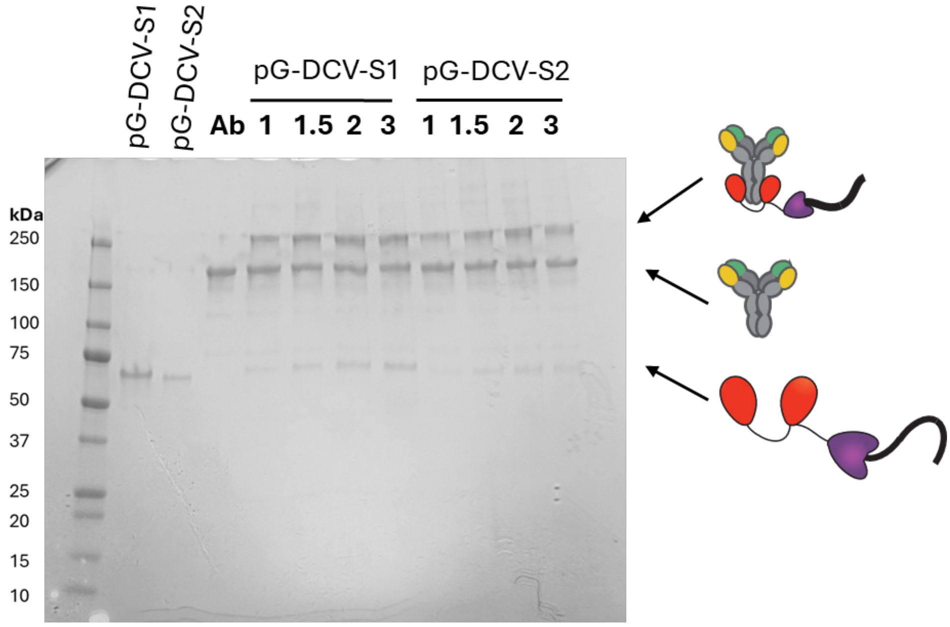


**Figure S5 SDS–PAGE showing the effect of pG–DCV–DNA excess over Adalimumab on PEA conjugates formation.** 1 µM of Adalimumab was reacted with 1/1.5/2/3 µM of pG-DCV-DNA (Table S4) and exposed to 365 nm UV light for 30 min. All samples showed partial photocrosslinking, with conjugate yield modestly increasing with the applied excess. Therefore, the next experiments were continued with a molar excess of 2 to maximize the complex formation and to minimize the amount of unconjugated pG-DCV-DNA.

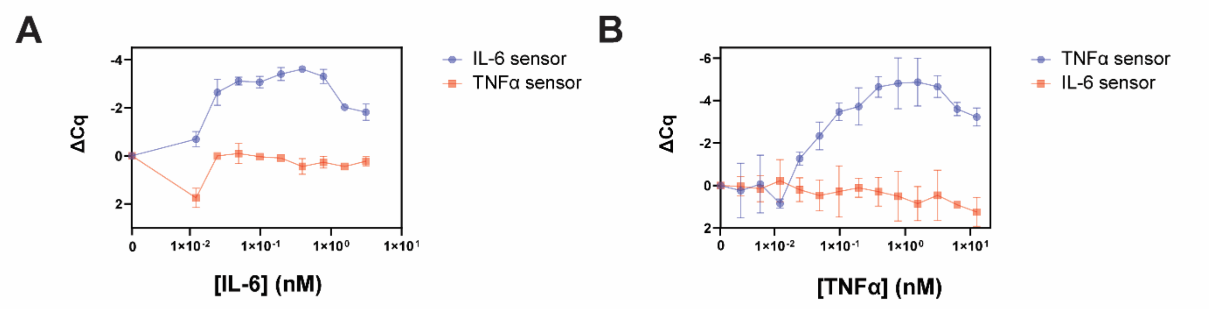


**Figure S6 PEA dose-response curves for Interleukin-6 (A) and TNFα (B).** 10 pM of the anti-IL-6 or Adalimumab conjugates were mixed with various concentrations of IL-6 or TNFα and incubated for 30 min at 37 °C. Next, T4 polymerase was added to the samples and the PEA reaction was performed for 90 min at 37 °C. The reaction mixtures were analysed by qPCR by mixing 1 µL of the reaction with 1x Taq polymerase MasterMix, EvaGreen® Plus dye and respective primers.
